# A dual sequence and culture-based survey of maize rhizosphere protists reveals dominant, plant-enriched, and culturable community members

**DOI:** 10.1101/2021.05.10.443483

**Authors:** Stephen J. Taerum, Jamie Micciulla, Gabrielle Corso, Blaire Steven, Daniel J. Gage, Lindsay R. Triplett

## Abstract

Protists play important roles in shaping the microbial community of the rhizosphere. However, there is still a limited understanding of how plants shape the protist community, and how well protist isolate collections might represent rhizosphere protist composition and function in downstream studies. We sought to determine whether maize roots select for a distinct protist community in the field, and whether the common or dominant members of that community are readily culturable using standard protist isolation methods. We sequenced 18S and 16S rRNA genes from the rhizospheres of maize grown in two sites, and isolated 103 protists into culture from the same roots. While field site had the greatest effect, rhizospheres in both sites had distinct protist composition from the bulk soils, and certain taxa were enriched in both sites. Enriched taxa were correlated to bacterial abundance patterns. The isolated protists represented six supergroups, and the majority corresponded to taxa found in the sequencing survey. Twenty-six isolates matched eight of the 89 core rhizosphere taxa. This study demonstrates that maize roots select for a distinct protist community, but also illustrate the potential challenges in understanding the function of the dominant protist groups in the rhizosphere.

**Originality-Significance Statement:** This is the first study comparing cultivation-dependent and independent methods for studying the protist community of plant roots, and the first untargeted analysis of the maize rhizosphere’s effect on protist communities. We show that maize in different sites select for distinct communities and overlapping enriched taxa, but that isolating the most important plant-associated protists may be a challenge for researchers.

## Introduction

Protists, or single-celled eukaryotes that are neither fungi, animals, nor plants, represent a significant part of the biomass and species diversity found in vegetated soils (Bardgett and Van Der Putten, 2014; Leach *et al*., 2017; Bass and del Campo, 2020). They include pathogens and commensals that that can impact plant health directly, through nutrient cycling, or through predation or stimulation of other microbes (Bass and del Campo, 2020). Despite the importance of protists, challenges posed by the extreme phylogenetic diversity and a lack of reference sequences has limited the exploration of protists in the plant microbiome. The challenge of isolating protists has similarly limited the scope of functional studies. As a result, understanding of the diversity and roles of protists in the plant microbiome lags behind that of bacteria and fungi.

Methodological advances have addressed some of the technical sequencing limitations for protists (reviewed in Keeling and Campo, 2017; and Geisen and Bonkowski, 2018), facilitating sequence-based exploration of protist diversity in plant environments. Studies have revealed that plant-associated protist communities typically consist of hundreds of species, the composition of which may be shaped by host species and tissue type, soil features, and management regimes (Dumack *et al*., 2020; Rossmann *et al*., 2020; Sun *et al*., 2021). Heterotrophic protists exert selective top-down influences in shaping the bacterial and fungal microbiome, and certain protist taxa may be correlated to disease development and other plant health indicators (Flues *et al*., 2017; Xiong *et al*., 2020; Sun *et al*., 2021). Studies in diverse species and soil environments are needed to determine whether these findings reflect universal patterns, and to understand the extent to which roots recruit protist communities, as they do bacteria and fungi (Sasse *et al*., 2018). Reports differ on whether rhizospheres select a distinct or reduced protist community from bulk soil (Sapp *et al*., 2018; Dumack *et al*., 2020), or whether there are differences in the soil compartments (Rüger *et al*., 2021; Sun *et al*., 2021). Very few studies have surveyed entire (i.e. untargeted) protist communities living in direct contact with plant surfaces, including the root-adhering soil compartment defined as the rhizosphere. We recently developed a PNA clamping approach to facilitate untargeted protist surveys in samples containing plant cells, by suppressing plant DNA amplification during 18S rRNA gene sequencing with universal primers (Taerum *et al*., 2020).

Mechanistic understanding of protist roles in plant health is largely informed by studies using cultured protist isolates. Addition of protist isolates to sterile plant soils revealed critical effects of selective predators on bacterial composition, metabolism, gene expression, and distribution, in ways that significantly affect plant health and biomass (Rosenberg *et al*., 2009; Jousset *et al*., 2010; Rubinstein *et al*., 2015; Weidner *et al*., 2017; Gao *et al*., 2019). One common method of isolating protists is the non-flooded petri dish method followed by serial dilution (Foissner, 1992; Howe *et al*., 2011): fresh soil samples are saturated with sterile water or buffer, then as microfauna emerge, droplets of the solution are repeatedly diluted and fed with heat-killed bacteria until a single protist morphotype is microscopically visible. The procedure requires time and skill, and protocols for long-term storage and distribution of the resulting isolates are limited. As a result, protist functional studies often employ in-house isolates or model species such as *Acanthamoeba castellanii* that are readily obtained from culture collections, but that have unclear relevance to the plant microbiome. As our taxonomic understanding of plant-associated protists develops, it will be important for the research community to develop isolation protocols and libraries targeted toward organisms of significance.

In this study, we performed a dual sequencing and isolation survey of protists in the maize rhizosphere in two research farms in Connecticut. We asked whether the root-attached soils of maize enrich for a distinct protist community from the immediately surrounding soil, and whether the dominant members of that community could be isolated in culture using the non-flooded petri dish and serial dilution method. 18S and 16S rRNA gene taxonomic profiling of rhizosphere soil revealed that protist and bacterial communities were significantly distinct from those of the immediately surrounding bulk soils, indicating plant enrichment of a rhizosphere protist microbiome, although the magnitude of compositional differences varied greatly between sites. We identified 89 core rhizosphere protist taxa, of which eight were both numerically abundant and significantly enriched in the maize rhizosphere, and found that these eight taxa correlated to patterns in the dominant bacterial community. We obtained and identified 103 protist isolates in pure culture from the same set of roots, and found that they generally represented taxa found in rhizosphere samples, including eight that were dominant and one that was enriched in the rhizosphere.

## Results

### Soil compartment and field site affect protist community composition

We sought to determine whether maize roots select for a distinct or reduced community of protists from the surrounding bulk soils. Taxonomic sequencing surveys were performed for maize fields in two sites in Connecticut, USA: one at the Griswold Research Center in Griswold, and the other at the Lockwood Farm in Hamden. Griswold soil is classified as a loamy sand, while Lockwood soil is a fine sandy loam; the sites also differed in planting history and weather exposure, and water content at sampling (Table S1). Soil that tightly adhered to maize roots upon shaking, or rhizosphere soil, was analyzed along with soil that was extracted along with the root crown but easily shaken off, defined here as bulk soil.

Bacteria amplicon sequence variants (ASVs) were classified to the genus level, while protist ASVs were classified to the species level. Animal and fungus ASVs were also recovered in the eukaryote dataset (Fig. S1) but were removed for the remainder of the analyses, along with plant and unclassified ASVs. The protist ASVs made up 27% of the total eukaryote dataset. Non-metric multidimensional scaling analysis of bacterial and protist sequences combined with PERMANOVA statistics revealed that bacterial, protist, and total eukaryotic communities significantly differed between sites (p < 0.001; Fig. 1A, Fig. S1A, Table S2). Rhizosphere and bulk soil compartments also had distinct bacteria and protist communities within sites (Table S2), but this difference was much more pronounced at the Lockwood site compared to the Griswold site (Fig. 1A, Fig. S1A), and with a greater level of significance at Lockwood (p = 0.008 vs 0.029 for bacteria, p = 0.009 vs 0.028 for the protists, and p = 0.007 vs 0.042 for the total eukaryotic community).

**Figure 1.**
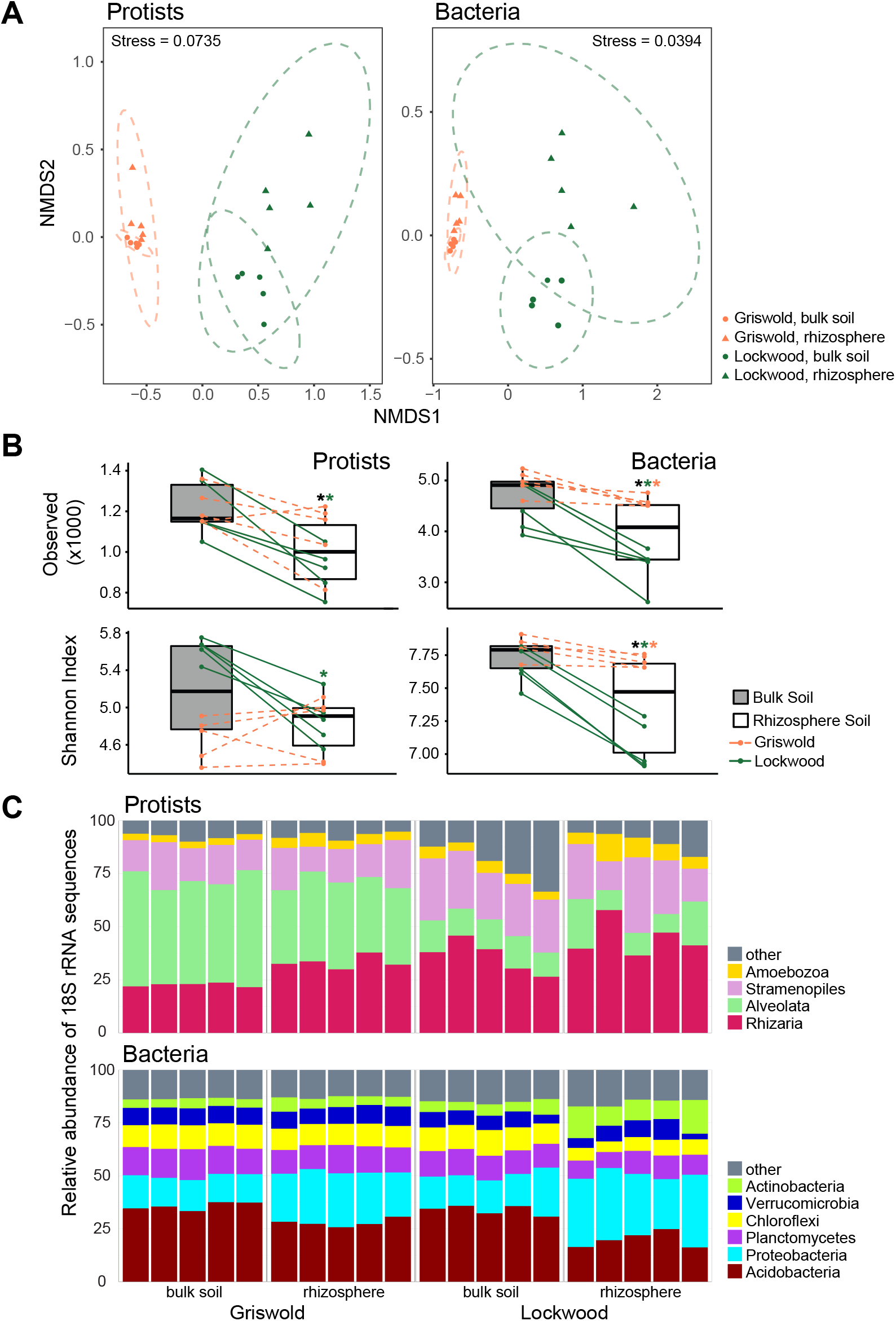
Community compositions of samples. **A)** NMDS plots showing the clustering of samples of eukaryotic microbes (animals, fungi and protists) and bacteria. Dashed-line ellipses indicate 95% confidence intervals. Stress values are indicated on the figure. **B)** Diversity of the protist and bacteria communities in the bulk soil and rhizosphere samples. Box plots indicate the median and quartiles of observed ASVs and Shannon’s diversity index, and points represent individual values. Lines connect points corresponding to the bulk and rhizosphere soil samples surrounding the same root. Green, orange, and black asterisks indicate statistical significance in Lockwood, Griswold, or across sites (p = 0.05). **C)** Relative abundances of the most dominant supergroup-level groups of protists and phylum-level groups of bacteria. The category “other” represents the sum of all other taxonomic groups.

### Rhizosphere effects on bacterial and protist community diversity vary by site

An emerging paradigm is that the rhizosphere selects for a bacterial community of reduced diversity relative to the bulk soil, and this has been demonstrated in the maize rhizosphere (Peiffer *et al*., 2013; Trivedi *et al*., 2020). We hypothesized that the same might be true for maize root protist communities. To test this hypothesis, we examined the effect of the soil compartment on bacterial and protist diversity. Bulk soils had similar levels of bacterial diversity and total eukaryotic diversity between sites, as measured by ASV richness and Shannon’s diversity index (Fig. 1B, Fig. S1B). In contrast, the bulk soil protist community had a significantly lower Shannon’s index in Griswold bulk soils than in Lockwood (p < 0.001) despite similar ASV richness, indicating a lower protist diversity at the Griswold site (Fig 1B). Bacterial diversity was significantly reduced in the rhizosphere compartments relative to surrounding bulk soil in both sites, with the difference more pronounced at Lockwood (Fig 1B). Protist community diversity at Lockwood was also lower in rhizosphere than in bulk soil (Fig 1B). There was no significant difference in protist diversity between compartments at the Griswold site, although there was a nonsignificant trend toward reduced ASV richness in the rhizosphere samples (p = 0.106, Fig 1B). These findings indicate that the maize rhizosphere can select for a reduced diversity of protists as it does for bacteria, but that the rhizosphere effect varies by site and is context-dependent.

### Maize rhizospheres were enriched in members of Rhizaria and Amoebozoa

We next analyzed proportional abundance of protist and bacterial taxa to identify broad taxonomic trends in the rhizosphere. Although bulk soils varied greatly in protist composition between sites (Fig. 1C), rhizosphere soils were significantly enriched in Rhizaria (p = 0.002) and Amoebozoa (p = 0.004) compared to bulk soils in both sites (Fig 1C, Fig S2, Table S3). Rhizospheres were also marked by a reduction in the dominance of Alveolata (p = 0.018). Examination of the five most abundant protist classes within each supergroup revealed that the Griswold protist communities were highly dominated by alveolates in the family Actinocephalidae, which comprised up to 45% and 34% of reads in Griswold bulk and rhizosphere samples, but only 1-3% of reads in Lockwood samples (Table S4). While several families showed differential relative abundance between soil compartments in at least one site, only the Allapsidae were among the dominant classes in the rhizospheres, but not bulk soils, in both sites (Table S4). Bacterial composition patterns also varied by site, but the rhizosphere was associated with an increased dominance of Protobacteria in both sites (Fig. 1C), consistent with previous studies (Peiffer *et al*., 2013).

### Identification of specific protist ASVs enriched in rhizospheres

Having identified compositional trends by supergroup, we next asked which protist ASVs were significantly more abundant in the bulk soil or rhizosphere compartments, and which protist ASVs differed in abundance between the two sites. Analysis using DESeq2 (Love *et al*., 2014) identified a small number of differentially abundant ASVs between soil compartments in each site (Fig 2A-B). Eleven ASVs were significantly enriched in the rhizosphere in Lockwood (Fig. 2A) and four were enriched in Griswold (Fig. 2B). All but one of the rhizosphere-enriched ASVs were in the supergroups Rhizaria or Amoebozoa, consistent with our earlier findings that these groups are enriched in relative abundance in maize rhizospheres (Fig. S2). Six of the fifteen rhizosphere-enriched ASVs were from the Rhizaria class Allapsidae, including four unique ASVs from the Group-Te genus, of which one ASV was enriched in both sites (Table S5). The other ASVs enriched in rhizospheres in both sites were amoeba of the LKM-74 lineage, also representing identical ASVs between sites, and two different ASVs of the genus *Hartmanella*.

**Figure 2.**
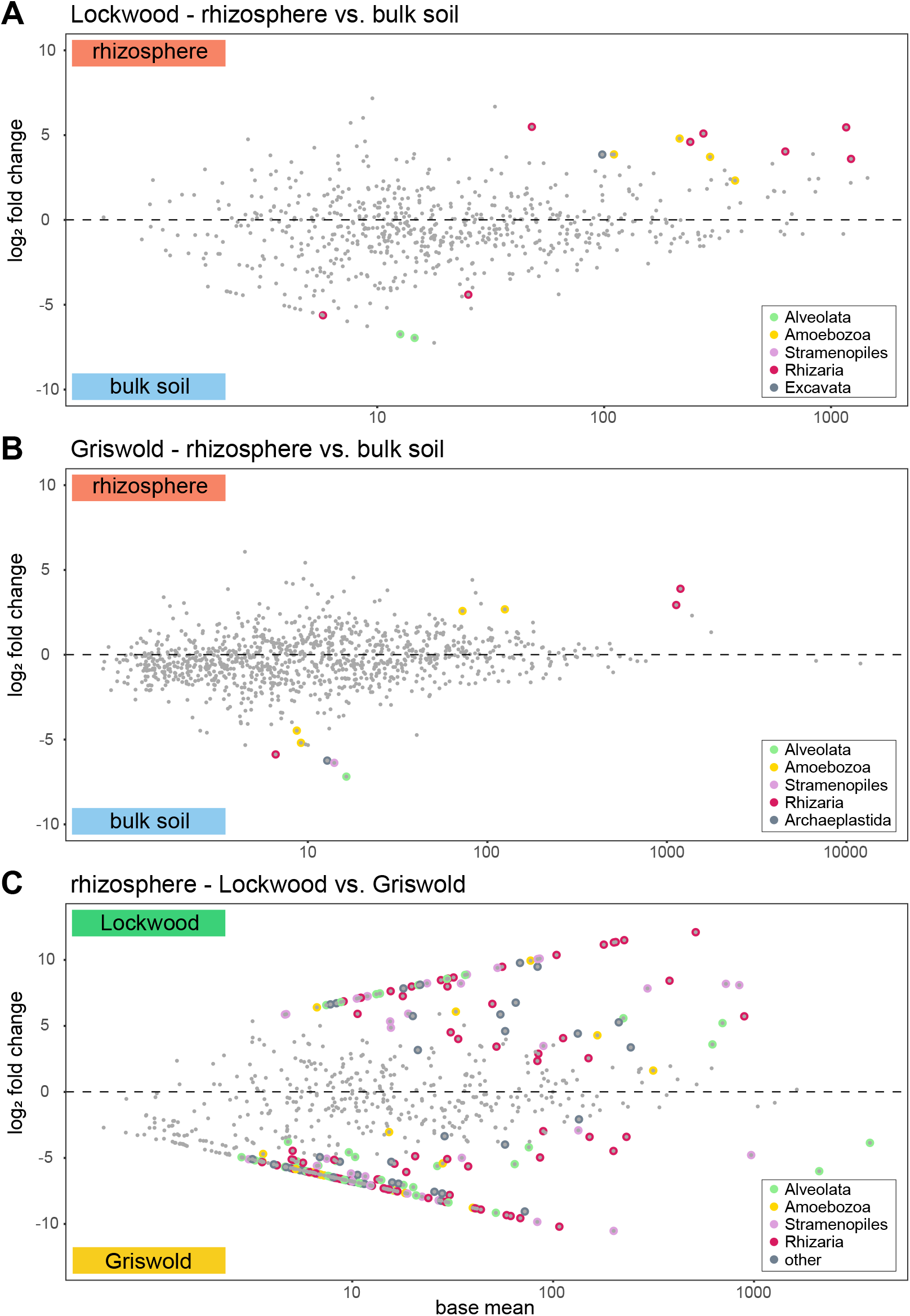
Differential abundances of protists between different treatments. **A-C)** MA plot based on DESeq2. Dots represent individual ASVs, with colors representing the supergroup designation of ASVs that are significantly more abundant in rhizosphere samples or bulk soil collected in Lockwood **A)** or Griswold **B). C)** MA plot showing the rhizosphere ASVs that are more abundant in Lockwood or Griswold. ASVs that are significantly more abundant in one environment are colored by supergroup. The category “other” represents the sum of all other taxonomic groups.

Four ASVs were enriched in Lockwood bulk soils and six in Griswold, representing five supergroups and with no phylum-level overlap between sites (Table S5). Despite a broader pattern toward enrichment of Rhizaria and Amoebozoa in the rhizosphere, half of the bulk soil-enriched taxa also belonged to these supergroups. We also identified over 200 taxonomically diverse ASVs that were differentially enriched between sites, which was expected given our previous finding that field site was the most important factor in driving protist compositional differences (Fig 2C). These results demonstrate that despite substantial site differences in bulk soil protist composition and soil compartment effects, maize rhizospheres enriched for an overlapping group of Rhizaria and Amoebozoa in both sites.

### Identification of core protist rhizosphere taxa across sites

Given the substantial variation in protist composition between soils, we next asked which protist ASVs were conserved in rhizospheres across sites. We identified the core protist taxa, which we defined as ASVs that were present in all ten rhizosphere samples. Of the 15755 ASVs detected across the rhizosphere samples, 89 were present in all ten rhizosphere samples, and eleven ASVs were represented with greater than 1% of reads in all rhizosphere samples (Fig. 3, Table S6). Half of the core taxa were in the supergroup Rhizaria, all of which represented the kingdom-level group Cercozoa (40 ASVs), while Stramenopiles, Ameobozoa, Alveolata, and Archaeplastida comprised 20, 14, and 9, and 6 core rhizosphere taxa, respectively (Table S6). The eleven core taxa representing >1% of reads included members of Amoebozoa, Cercozoa, the alveolate class Colpodida, and the plant parasitic oomycete order Peronosporales (Fig. 3, Table S6). The core taxa also included eight of the thirteen ASVs found to be significantly enriched in the rhizosphere in one or both sites, including the Amoebozoa LKM-74 and the Cercozoa Group-Te lineages. In addition to Peronospora, other core rhizosphere clades associated with plant parasitism included Spongospora and multiple Pythium ASVs.

**Figure 3.**
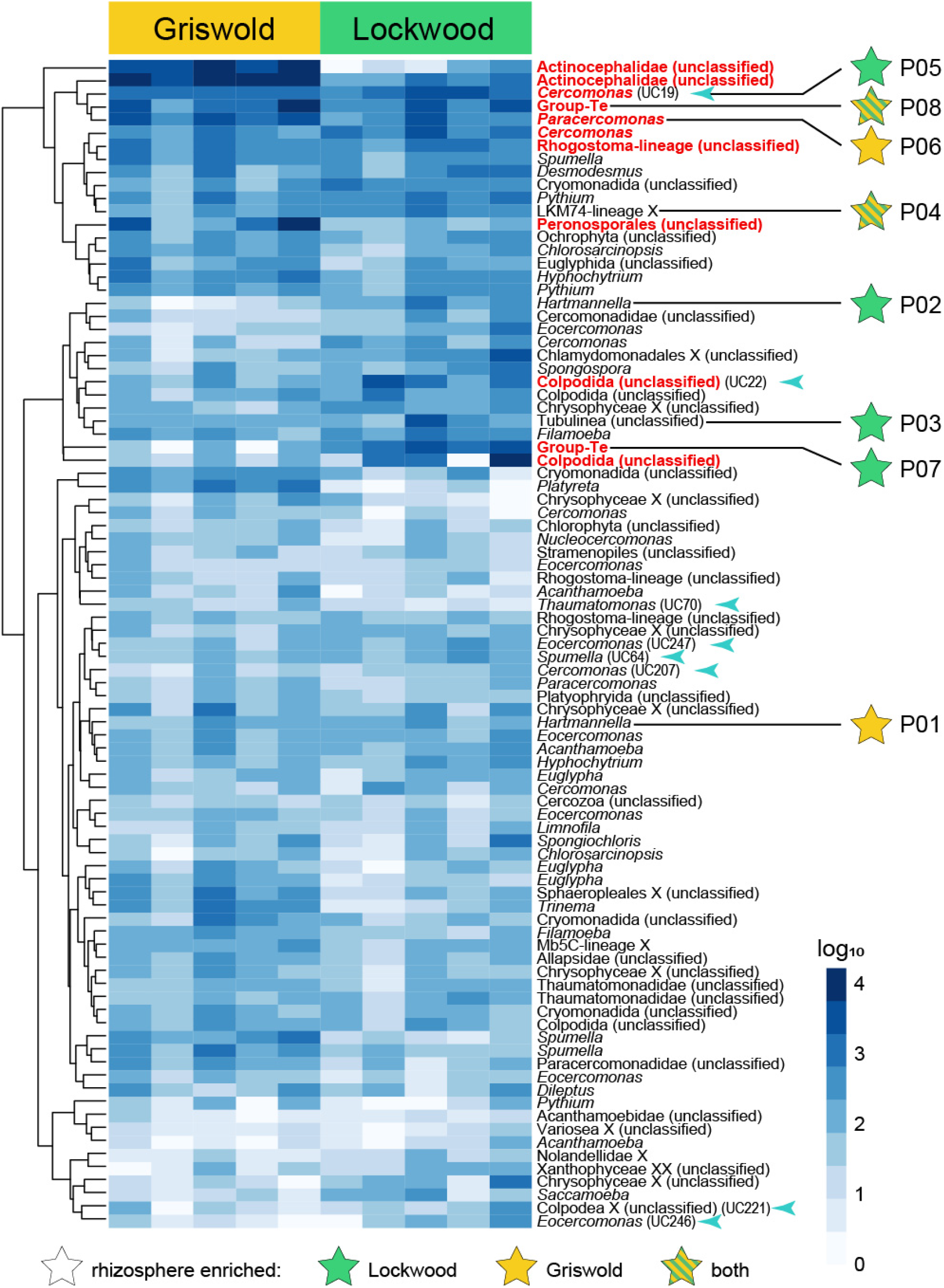
Core protist taxa. Heatmap showing the abundance in log_10_ scale of the 89 protist ASVs that are present in all ten rhizosphere samples. ASVs are vertically arranged according to the dendrogram, which indicates similarities in relative abundances across samples. The 11 ASVs that had a relative abundance of 1% or higher are shown in red text. Stars indicate ASVs that were enriched in the rhizosphere in one or both locations, based on the DESeq2 analysis. Arrowheads and culture codes indicate ASVs that matched the V9 regions of cultures that were isolated in this study.

### Correlative relationships between abundant protists and bacteria in the maize rhizosphere

Protist community composition can be shaped by soil bacteria in addition to the abiotic and host environmental factors, but protists can also exert top-down selection of bacterial communities through predation or symbiosis (Rossmann *et al*., 2020). We employed canonical analysis of principal coordinates to identify bacterial populations that may have explanatory patterns in protist compositional data. The 15 most abundant bacterial taxa across all 20 samples and the composition of the different protist communities were interrogated. Proteobacteria in the genera *Massilia, Pseudoduganella*, and *Sphingomonas* sp. were associated with the Lockwood rhizospheres, along with a Verrucomicrobia from the family Pedospheraceae and a member of the class Anaerolinae (Fig. 4, bacterial taxa listed in Fig. 5). A Rokubacteria sp. influenced the community toward Lockwood bulk soils. The Griswold protist communities, while not greatly distinct by compartment, were predominantly influenced by six Acidobacteria including three Pyrinomonads (Fig. 4).

**Figure 4.**
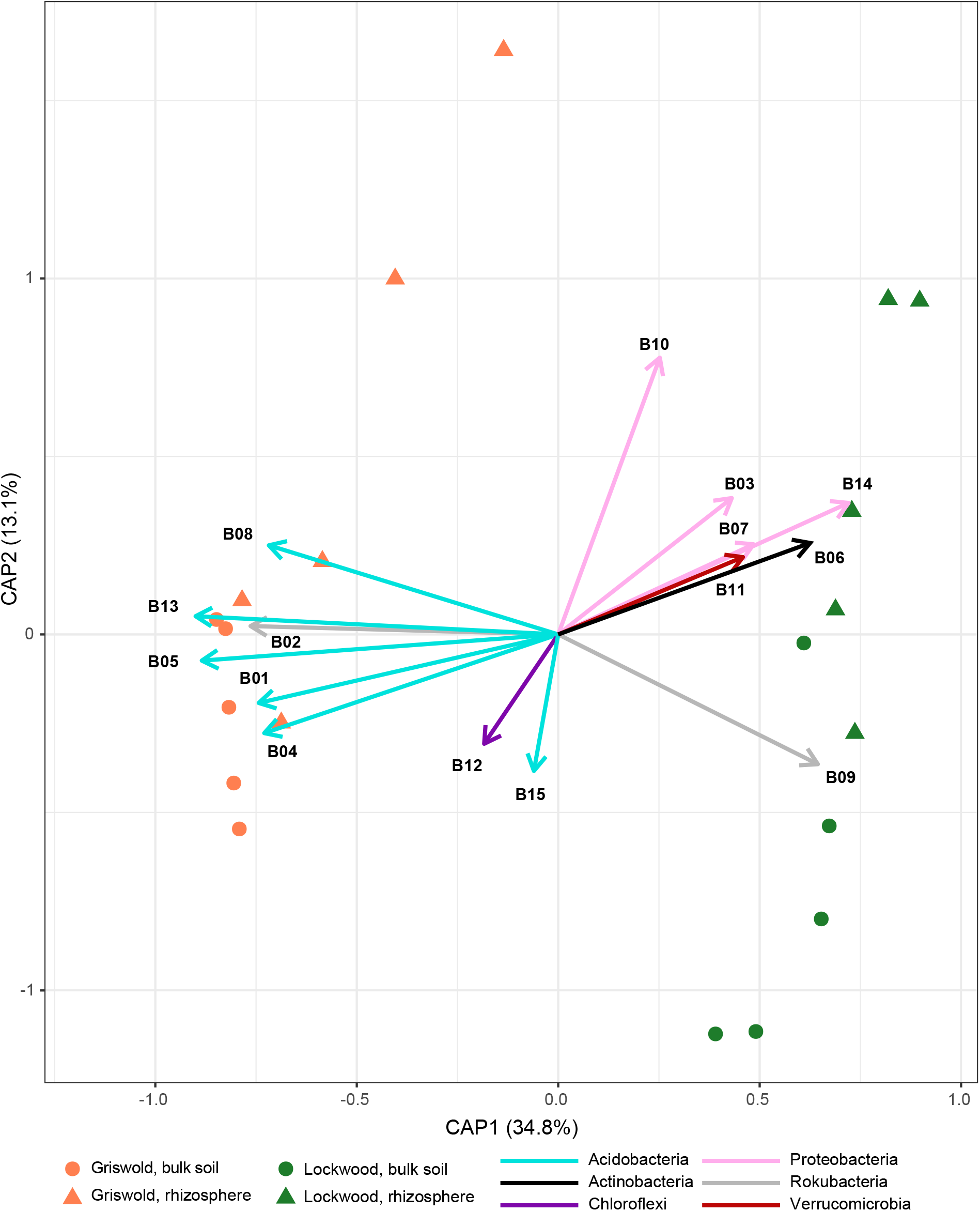
Canonical analysis of principal coordinates of the protist ASVs. The first two principal coordinates are shown on the axes. The vectors indicate the impact of the 15 most abundant bacterial ASVs on the observed variance in the plot. The vector directions indicate how a bacterial ASV is impacting the protist community, and the vector lengths indicate the magnitude of the impact. The bacterial vectors are colored based on their phylum.

**Figure 5.**
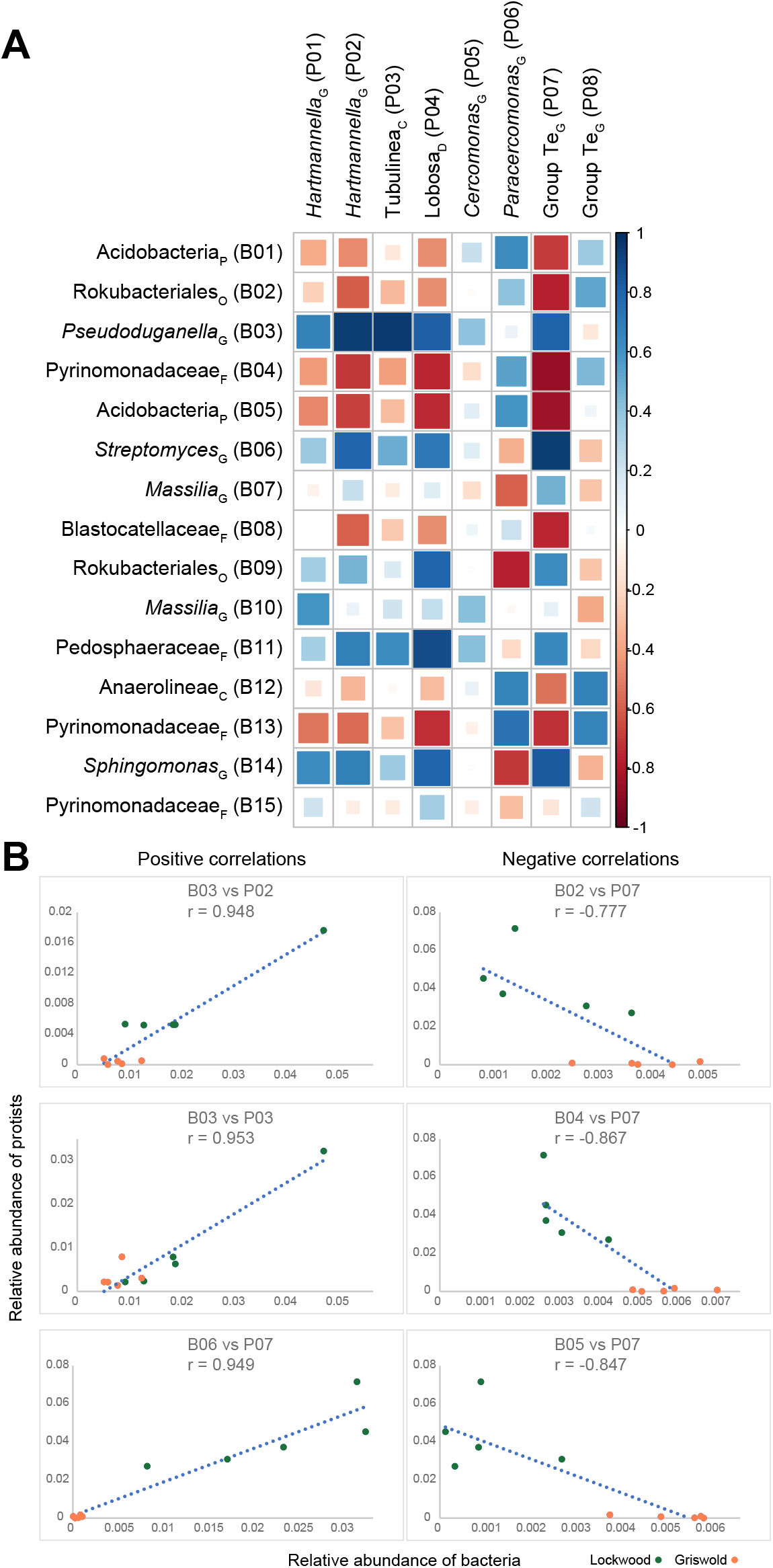
Correlations between the abundances of bacteria and protists. **A)** Correlogram indicating the Pearson’s correlation r-values between the relative abundances of the eight protist ASVs that were enriched in the rhizosphere samples (based on DESeq2), and the 15 most abundant bacterial ASVs. Colors and sizes of the squares indicate the strength and direction of the correlations. **B)** Dot plots with trend lines of the three strongest positive and negative correlations between protist and bacterial relative abundances. r-values are indicated for each association.

Pearson’s correlation analysis was used to identify co-occurrence relationships between relative abundance of the numerically dominant bacteria and abundance of the eight core, rhizosphere-enriched protist ASVs listed in Fig. 3. The four Amoebozoa ASVs were positively correlated with the abundance of *Pseudoduganella, Streptomyces*, and *Sphingomonas* sp., and were negatively correlated with several members of the Acidobacteria (Fig. 5). Cercozoa exhibited more variable correlations: P08, the Group-Te ASV that was rhizosphere-enriched in both sites, was positively correlated with several Acidobacteria, while another Group-Te ASV (P07) exhibited similar correlative patterns as the Amoebozoa. These data demonstrate that abundant bacteria are associated with both field site and soil compartment differences in protist community composition, and identify specific co-occurrence patterns between abundant bacteria and protists.

### Isolation and identification of rhizosphere protists

Metataxonomic sequencing is highly useful for profiling microbial communities and identifying ecological patterns but isolates or defined communities will be critical for downstream testing of mechanistic hypotheses. We predicted that a library of rhizosphere protists isolated using standard methods would include some taxonomic overlap with the 18S rRNA gene sequencing results, although it would likely be missing groups that are unculturable or may contain taxa that are not amplified by V9 primers. Protists were grown and isolated from root samples of the same ten plants used for 16S/18S rRNA gene sequencing, using serial dilution heat-killed *Escherichia coli* as a food source. Amoeboid protists were not selected for isolation, as we wanted to avoid culturing any of the abundant amoebae that are opportunistic human pathogens or disease vectors (Marciano-Cabral and Cabral, 2003; Balczun and Scheid, 2017). In addition, experimenter decision making factored into isolate selection; protists were not selected for continued culture once several morphologically identical protists had already been cultured from the site. A collection of 250 protists were initially isolated, and of these 103 survived multiple serial passages in culture and were sequenced. Partial 18S sequencing revealed that 101 of the isolates contained 52 unique V9 sequence groups (Table 1, Table S7). We obtained only partial V9 sequence for the other two cultures. Although isolation results are not quantitative, the unique isolates predominantly consisted of Stramenophiles (22), Rhizaria (16), and Alveolatea (9), with two isolates of Haptista (also referred to as Hacrobia) related to *Chlamydaster* sp., and single isolates of an Opisthokonta related to the class Nucleariidea, an Excavata in the genus *Stachyamoeba*, and an amoeba related to *Flamella sp*. Thirty-six unique isolates (71%) had V9 regions matching ASVs detected in 18S rRNA gene sequencing studies. Thirty-three isolates matched ASVs sequenced from rhizosphere samples, and eight isolates matched a member of the 89 core ASVs, with one isolate (a species of *Cercomonas*) matching an ASV that was enriched in the rhizosphere in Lockwood (Fig. 3). An additional unique isolate (UC43) was present in all rhizosphere samples, but was not considered to be one of the core protists because the matching ASV was classified as “unclassified eukaryote” by the PR^2^ database. The eleven unique isolates not representative of sequencing data included two members of the supergroup Haptista (also referred to as Hacrobia). In summary, the isolation survey captured taxa that included diverse representatives of the sequencing survey as well as some taxa absent from the sequencing, and were representative, at the ASV-level, of several of the dominant and rhizosphere-enriched taxa identified by high-throughput sequencing.

**Table 1.**
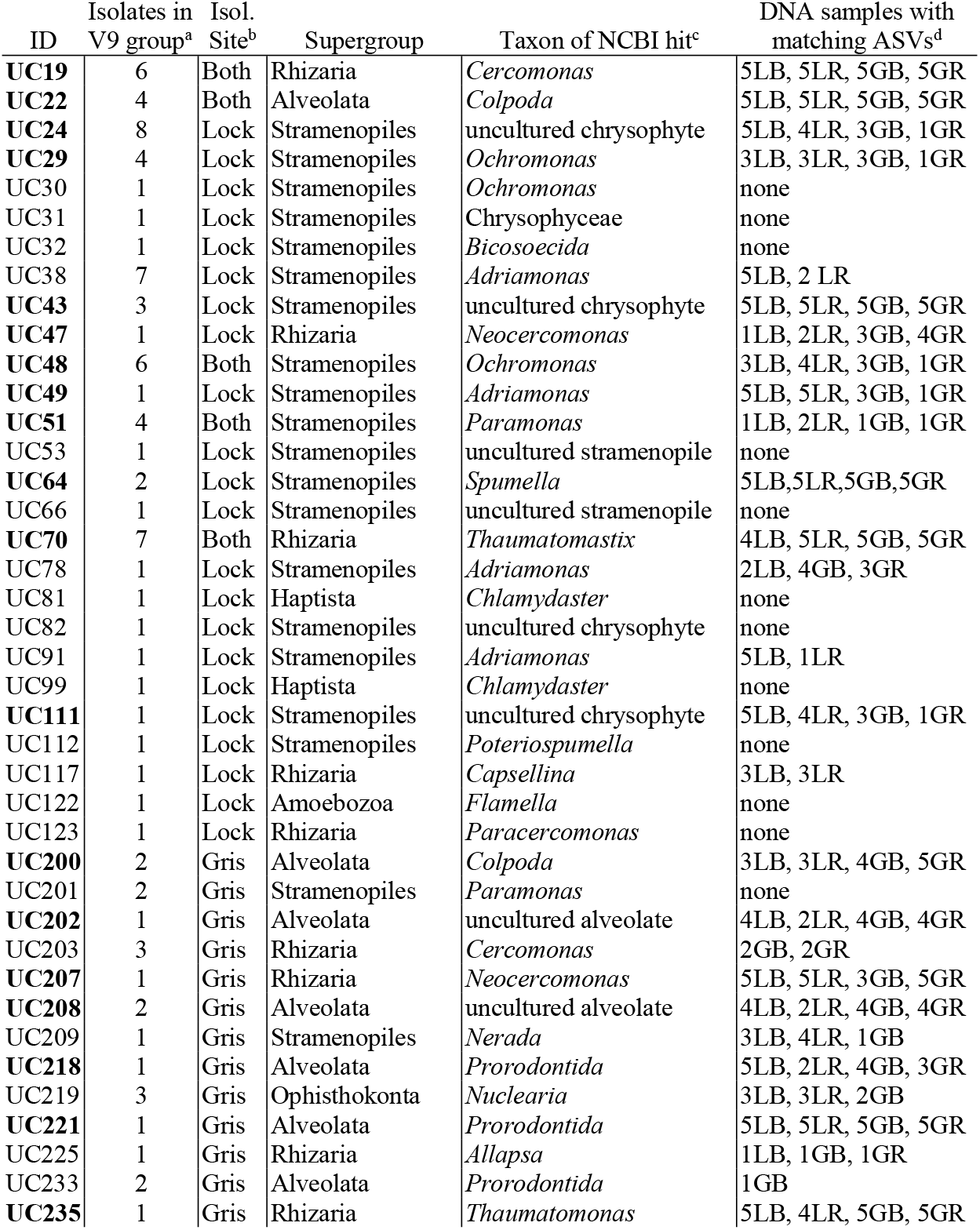
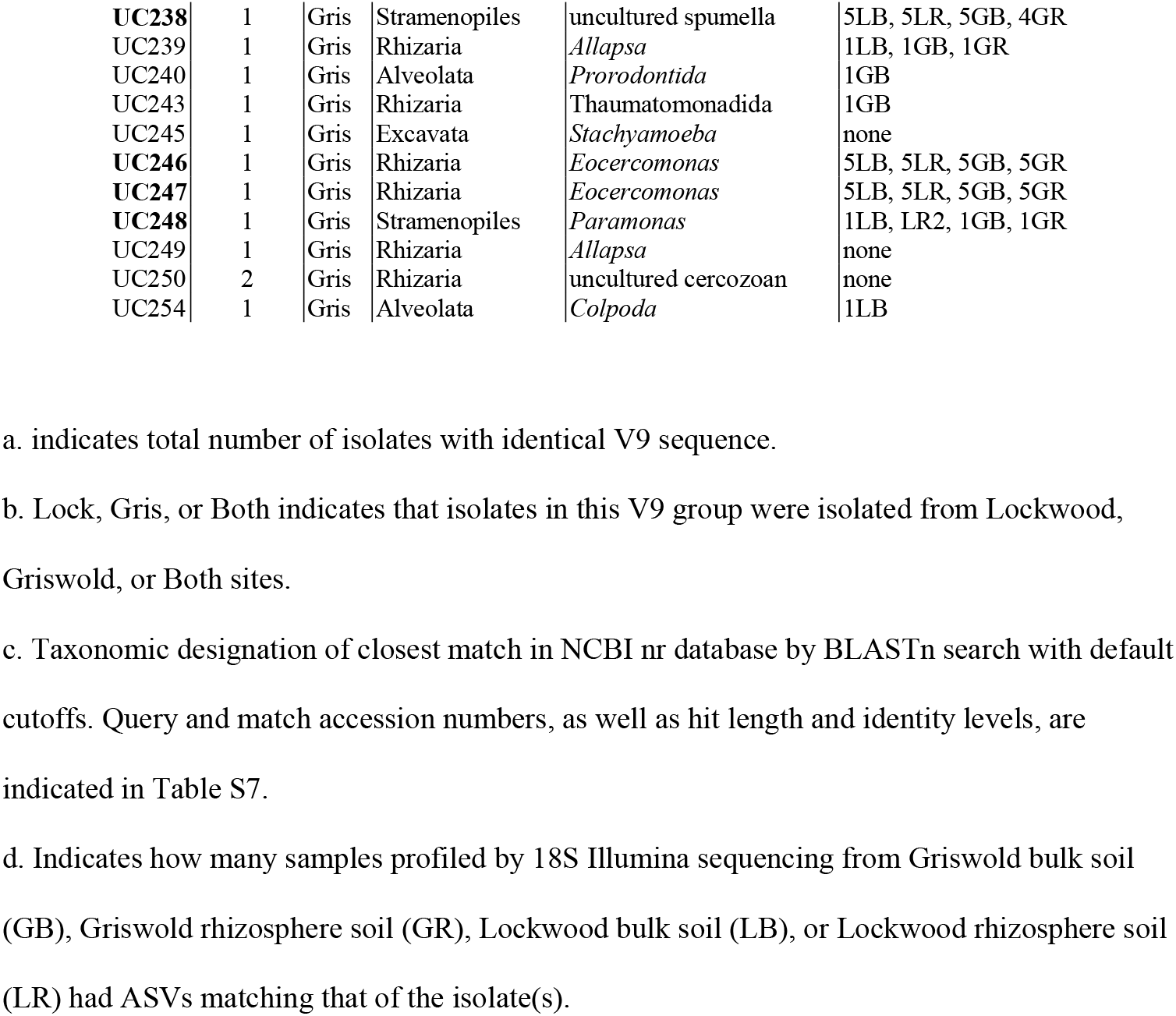
Protists isolated in culture from maize rhizospheres. Number of isolates with identical V9 sequence, isolation site(s), predicted classification, and samples with matching V9 sequence in 18S taxonomic profiling study are shown. Bold numbers indicate isolates with ASVs matching those sequenced in rhizospheres in both sites.

## Discussion

Linking protist traits to ecological function is a major goal toward optimizing the microbiome for plant health (Gao *et al*., 2019), and developing isolate resources will be a critical element in reaching that goal. This study investigated maize rhizosphere effects on the protist community, and surveyed the cultivable portion of that community as a step toward linking direct and ecological understanding. We found that across sites with differing soil properties and microbiota, the maize root surface hosts a distinct protist community from the immediately surrounding soil, enriching for overlapping clades within the Rhizaria and Amoebozoa, and that the abundance of rhizosphere-enriched protists are correlated to that of dominant bacteria. The isolation survey generated a diverse protist library predominantly representing taxa found in the sequencing survey, including some that were dominant in that survey.

We observed that soil compartment affected the maize protist community at both experimental sites, but substantial differences in protist composition and diversity were only found at the Lockwood site. The two sites differed in soil type, climate, planting history, and bulk soil microbial communities. The most notable difference was the extreme dominance in Griswold of the Apicomplexa clade Actinocephalidae, known as host-specific parasites of animals that dominate rainforest protist communities (Mahé *et al*., 2017), but which are not nearly as dominant as Cercozoa in most other soils (Oliverio *et al*., 2020). It is not known why a temperate agricultural field might be dominated by Actinocephalidae, but the finding that the dominance was reduced in the rhizosphere indicates that they were not associated with a maize pest. The Lockwood site had the clearest bulk-rhizosphere differences in diversity and composition of bacteria, fungi, and metazoans (Table S2), suggesting that factors affecting the strength of the rhizosphere effect of protists and other microbes may be linked. It is possible that the drier, sandier soil at Griswold affected the root influence into the non-attached soil we defined as bulk, and that bulk sampling further from the plant may have revealed larger differences. Other factors such as plant age, climate, and soil management history have been previously found to influence the strength of the rhizosphere effect on bacterial microbiota (Berg and Smalla, 2009; Philippot *et al*., 2013; Nuccio *et al*., 2016).

Despite the large site-specific differences in protist composition, Rhizaria and Amoebozoa were significantly enriched in maize rhizospheres, while Alveolata were depleted. ASVs enriched in the rhizospheres were associated with Lobose amoebae or Cercozoa, with the uncultured genus-level clade Group Te (family Allapsidae) representing a third of the enriched ASVs. Intriguingly, Group Te protists were previously found to comprise five of twelve taxa enriched in the rhizospheres of *Arabidopsis* potted in German soils, and were also observed to form a major network hub in the potato rhizosphere (Sapp *et al*., 2018; Dumack *et al*., 2020). Another *Allapsa* sp. was among the taxa most predictive of tomato health in the presence of bacterial disease pressure (Xiong *et al*., 2020). The finding that Group Te is also enriched in monocot rhizospheres from two North American soil communities suggests that it is a plant-associated clade of widespread significance. Other taxa enriched in both sites included *Hartmanella vermiformis* and LKM74-group (Dermamoebidae) amoebae, both of which are widely distributed in soils (Takenouchi *et al*., 2016; Blandenier *et al*., 2017). *Hartmanella* sp. were previously observed to be enriched in a maize rhizosphere soil microcosm collected from a field in Germany (Zhang and Lueders, 2017), but LKM74-group amoebae have not been previously documented as rhizosphere-enriched to our knowledge.

Some of the enriched protist taxa were correlated to the abundance of common bacterial groups. For example, the *Hartmanella* and LKM74-group from both sites were negatively correlated to Acidobacteria including Pyrinomonadaceae, but positively correlated to the Verrucomicrobia family *Pederosphaeraceae*, the Burkholderiales genus *Pseudoduganella*, and *Sphingomonas* and *Streptomyces* sp. Correlations could arise from similar environmental pressures and are not an indicator of mechanism, but may be useful for forming hypotheses for future study of protist-bacterial dynamics. *Verrucomicrobia* and *Duganella-*like sp. are common protist symbionts (Sato *et al*., 2014; Khojandi *et al*., 2019), and *Sphingomonas* and *Streptomyces* sp. have been identified as predation-resistant, likely due to secondary metabolite and spore forming capabilities (Pérez *et al*., 2011; Zou *et al*., 2020).

A major goal of this study was to generate a cultured isolate library from sequence characterized samples, as a step toward unifying sequence-based understanding with organismal biology. Isolation surveys are limited to free-living organisms, and our study was further limited to non-amoeboid bacterivores. The isolate library nevertheless represents a variety of rhizosphere protist taxa. Some of these protists are representatives of soil-inhabiting genera isolated in previous targeted studies, such as *Thaumatomastix, Paracercomonas*, and *Eocercomonas* (Bass *et al*., 2009; Howe *et al*., 2011), while others have no close homolog among characterized cultures. Further morphological and trait characterization of the isolates is underway with the objective of their deposition in culture collections; because many of our isolates are from genera with no reference genomes, they may contribute to the sparse genomic resources for for Ochrophyta, lobose amoebae, and especially Rhizaria (Sibbald and Archibald, 2017). Several protists that were isolated and cultured in our study did not have matching ASVs in any sequenced samples, possibly due to their rareness before culturing, or because the primers used for the high throughput amplicon sequencing may have not amplified certain groups (Vaulot *et al*., 2020). Several isolates were cultures that were “core” rhizosphere taxa, or closely related to those abundant or rhizosphere-enriched protists identified by high-throughput sequencing, and thus might serve as a valuable resource for study of the contribution of certain taxa or lifestyles to the plant microbiome. A caveat is that in some cases different protists from the same genus were enriched in different soil compartments, and previous work has established that even closely related species can differ in grazing patterns and bacterial co-occurrence patterns (Flues *et al*., 2017). Interspecies variation should be taken into consideration when developing experimental studies from available cultures.

Although some protists of interest have not been isolated yet, taxonomic survey data can be valuable in prioritizing future isolation targets for plant health research, especially in guiding decision making regarding the sampling depth of common morphologies. For example, our depth of sampling within Rhizaria may have been more comprehensive if we had pre-selected common morphotypes for deep isolation and culturing. Study of organisms related to previously uncultured targets, such as the Group Te relative *Terretomonas rotunda* (Howe *et al*., 2009) could also provide useful insight to morphotypes and nutritional needs of the target species. For example, the first isolate of a LKM74-clade amoeba was reported in 2016, and was found to form unique visible structures that could be targeted in isolation by dilution or micromanipulator (Blandenier *et al*., 2017). Single-cell transcriptomes and metatranscriptomes technologies could soon help reveal protist nutritional needs, but classic growth requirement studies are still highly valuable; dedicated survival studies of the marine flagellate *Oxyrrhis marina* led to the complex media that helped establish this organism as a model system in tidepool ecology (Lowe *et al*., 2011). Network analyses might help identify preferred prey or nutritional symbionts to enrich for desired protist groups. This study demonstrates the feasibility and value of pairing culture-dependent and independent descriptions of protist diversity, toward future integration of ecological patterns with functional traits and molecular mechanisms.

### Experimental Procedures

#### Collections

Rhizosphere sample collection was performed as described in Taerum *et al*. 2020; rhizosphere soil samples L1, L2, G1, and G2 used for clamp technique validation in that paper are synonymous with samples Lrh1, Lrh2, Grh1, and Grh2 from this work (i.e., the same soil and DNA samples were used for those four samples across two papers, but separate library preparations and sequencing runs were performed for this study). Samples were collected from plots of B73 maize planted May 15^th^ and 17^th^, 2019 in Griswold and Hamden, Connecticut, USA. Rainfall and temperature data were recorded at weather stations within 100 yards of each field. Soil characteristics and sampling metadata are listed in supplemental Table S1. Root crowns were harvested from 5 plants per plot, selected randomly from different rows, eight weeks after planting on July 10^th^ and 12^th^ (V6 stage). Root crowns were removed carefully from soil with a clean spade so as to retain most of the soil inside the top part of the root crown. Root crowns were then shaken vigorously for one minute to remove loosely adhering soil, which was collected into an autoclaved pan. This loosely adhering soil was then mixed well and sampled as bulk soil. From each plant, five roots were selected at random from among the crown and seminal roots, and these were collected and trimmed to 10 cm below the soil line with sterile scissors. The root samples were placed in 35 mL cold sterile phosphate buffered saline and vortexed for 2 minutes, following the protocol of McPherson et al. (2018). Samples were immediately placed on ice and placed in a -80°C freezer until DNA extraction. The remaining portion of the root crown was stored on ice in clean plastic bags and refrigerated until protist isolation could be performed.

### DNA extraction, amplification and sequencing

Rhizosphere and bulk soil DNA was extracted using the DNeasy PowerSoil Kit (Qiagen, Germantown, MD, U.S.A.), following the manufacturer’s protocols. DNA was stored at -80C until amplification.

Bacteria and eukaryote libraries were generated for all ten rhizosphere samples and all ten bulk soil samples in single PCR reactions. We amplified and sequenced two technical replicates per sample of each bacteria and eukaryote library to ensure that there were minimal differences in the amplified communities between the replicates. After confirming this, we merged the two replicates. Bacterial libraries were amplified using the primers 515F (5’-GTGYCAGCMGCCGCGGTA-3’) and 806R (5’-GGACTACVSGGGTATCTAAT-3’), which targeted the V4 region of the 16S rRNA gene. Eukaryotic libraries were amplified using the primers 1391F (5’-GTACACACCGCCCGTC-3’) and EukBr (5’-TGATCCTTCTGCAGGTTCACCTAC-3’), which targeted the V9 region of the 18S rRNA. The primers were attached to Illumina adapters and indices. Each 25 µL reaction contained 1 U Invitrogen Platinum *Pfx* DNA polymerase (Thermo Fisher), 1x reaction buffer, 0.2 mM dNTPs, 1.5 nM MgSO_4_, 0.5 µM each primer, and 0.2 ng/uL template. The eukaryotic reactions also contained 3.75 µM of the PNA clamp PoacV9_01 (Taerum *et al*., 2020) to suppress amplification of plant DNA. PCR amplifications consisted of a denaturation step of 3 min at 95°C, followed by 30 cycles of a denaturation step of 30 s at 95°C, an annealing step of 30 s at 55°C, and an extension step of 30 s at 72°C; followed by a final extension step of 5 min at 72°C.

Libraries were cleaned and normalized to 1 ng/µL using the SequalPrep™ Normalization Plate Kit (Thermo Fisher), following the manufacturer’s directions. The bacterial and eukaryotic libraries were pooled separately. Each pooled library was sequenced using 2 x 250 bp chemistry on the Illumina MiSeq platform.

### Bioinformatics pipeline

#### Assembly, filtering, and classification

Sequence reads were assembled and filtered using mothur v. 1.44.0 (Schloss *et al*., 2009). Bacterial contigs were filtered to have a maximum length of 300 bp in length, while containing no ambiguous bases, and a maximum homopolymer length of eight. Eukaryotic contigs were filtered to be between 90-250 bp in length, while containing no ambiguous bases, and a maximum homopolymer length of eight. Chimeras were identified using VSEARCH (Rognes *et al*., 2016), and removed using mothur. Representative bacterial amplicon sequence variants (ASVs) were classified against the SILVA v. 132 database (Quast *et al*., 2013), while representative eukaryotic ASVs were classified against the Protist Ribosomal Reference (PR^2^) database (Guillou *et al*., 2013). Classification was done using the naïve Bayesian classifier in mothur (Wang *et al*., 2007), with a bootstrap cutoff of 80%.

#### Statistics and composition

Data were imported into the phyloseq package (McMurdie and Holmes, 2013) which was implemented in R. ASVs were counted in both datasets. α-diversity statistics were calculated after subsetting the datasets to the samples with the smallest numbers of reads. Pairwise differences in ASV richness and Shannon’s diversity were analyzed using a Student’s t-test for paired samples. Permutational multivariate analysis of variance (PERMANOVA) tests were conducted based on Bray-Curtis similarity matrices to determine if there were differences in bacteria and eukaryote composition between the rhizosphere and bulk soil samples, as well as between the sample locations. Homogeneity of dispersion was calculated to determine if the treatments had homogenous variances. Differences in community composition were visualized using non-metric multidimensional scaling (NMDS) plots.

#### Differentially abundant ASVs

We used the DESeq2 package (Love *et al*., 2014) implemented in R to determine the eukaryotic ASVs that were enriched in the rhizosphere compared with the bulk soil, as well as the eukaryotic ASVs that were differentially abundant in the two locations. DESeq2 fits a negative binomial test to each ASV and uses a Wald test to compare between groups. ASVs that were present in fewer than three of ten rhizosphere samples were removed to prevent ASVs that are highly abundant in one sample from creating spuriously significant differences. The ASVs were then visualized on a MA plot.

#### Core protist taxa

To determine the core taxa within the rhizosphere communities, we removed the protist ASVs that did not occur in all ten rhizosphere samples. The remaining 89 ASVs were then plotted on a heatmap using the R package microbiomeutilities (Shetty and Lahti, 2018). To determine the most common core ASVs, the samples were again filtered so that only ASVs that had an abundance of 1% or greater were retained.

#### Co-occurrence with bacteria

Canonical analysis of principal coordinates (CAP) was conducted to identify bacterial taxa that may influence the composition of the eukaryote communities. The relative abundances of the bacterial ASVs were calculated in Phyloseq, after which the 15 most abundant bacterial ASVs in all combined reads were selected. The bacterial relative abundances were incorporated into the eukaryote metadata, after which CAP was conducted in phyloseq. The bacterial ASVs were plotted as vectors on the CAP plot.

### Protist isolation and 18S identification

#### Protist Isolation and Culturing

25g of root tissue was removed from the root ball and transferred to a sterile petri dish containing 5mL of sterile water. Dishes were gently shaken to coat all root tissue with water. Dishes were sealed with parafilm and stored in the dark at room temperature for 48 hours. Rhizosphere suspensions were diluted until 1µL droplets contained single protists. These were transferred to individual wells in a 96-well microtiter dish containing 100µL Page’s saline solution + 10% soil extract solution (SES; the soil used to make this solution was from Lockwood farm) with heat-killed *E. coli* DH5α (OD_595_ = 0.005) as a food source (Page, 1976). The microtiter dish was kept in the dark at room temperature for one week, allowing for the individual protists to reproduce. Uniform protist cultures were transferred to Nunc Cell Culture Tubes (ThermoFisher #146183) containing 1.5mL Page’s Saline Solution + 10% Lockwood SES with heat-killed *E. coli* DH5α (OD_595_ = 0.005) as a food source for long term storage.

#### Protist Sequencing

Protists were displaced from the bottom of 25 cm^2^ culture flasks containing 10 ml of Page’s saline solution + SES and heat-killed *E. coli* with a sterile cell scraper. 1mL of protist culture was transferred to a 1.5mL tube and centrifuged at 13,000rpm for 5 minutes. The protist cell pellet was resuspended in 50uL of supernatant and used in the Wizard Genomic DNA Purification Kit (Promega #A1120). gDNA was quantified using a NanoDrop. In some cases, protist culture was used directly for amplification of 18S rDNA. A 20µL PCR was set up as follows: 1X GoTaq Green Master Mix (Promega #M7122), 50ng gDNA, 0.5µM 18S_FU primer (ATGCTTGTCTCAAAGGRYTAAGCCATGC), 0.5µM EukBR primer (TGATCCTTCTGCAGGTTCACCTAC), nuclease-free water to 20µL. Reactions were run on a thermocycler at the following settings: 95°C for 2 minutes, 35 cycles of 95°C for 1 minute, 55°C for 1 minute, 72°C for 90 seconds, and a final step of 72°C for 5 minutes. After amplification, the PCR products were purified using the Wizard SV Gel and PCR Clean-Up System (Promega #A9281). Protist 18S PCR products were quantified using a NanoDrop, then diluted to 5ng/µL and sequenced with both 1209F (CAGGTCTGTGATGCCC) and EukBR primers at Genewiz (New Jersey, USA). Forward and Reverse sequencing reads were assembled into contigs, then run against the NCBI Nucleotide Blast Database.

## Data Availability

All Illumina and Sanger sequences generated for this project can be found in NCBI GenBank under BioSample PRJNA714637 and accession nos. MW775149-251. Protist isolates are available upon request from the authors with the appropriate permits; work is underway to optimize long-term storage methods for deposition of the isolates into public culture collections.

## Acknowledgements

This work was supported by AFRI Foundational Program grant from the United States Department of Agriculture-National Institute of Food and Agriculture (USDA-NIFA) to L.R. Triplett, D.J. Gage, and B. Steven (grant 2019-67013-24412). Kate Manning and Carlos Calderon provided field assistance and were supported through an AFRI Education and Learning Initiative grant from USDA-NIFA (grant number 2017-67032-26013). We thank Ms. Regan Huntley and Ms. Kavya Uddaraju for technical lab assistance.

## Supplementary Data

**Table S2** PERMANOVA and dispersion statistics for comparisons between sites and between treatments. Statistics are shown for bacteria and all eukaryotes combined, as well as protists, animals and fungi separately.

**Table S3** Two-way analysis of variance (ANOVA) of the relative abundances of four most abundant protist supergroups across all samples. Site and soil compartment are the treatment factors.

**Table S4** Relative abundances of the five most abundant families within each supergroup for each site and compartmental group. Taxa representing greater than 5% of all reads are indicated in bold text.

**Table S5** Differential abundance statistics from the DESeq2 analyses used to generate the MA plots in Fig. 2. The table is colored according to the compartment (blue = bulk soil, red = rhizosphere) or site (orange = Griswold, green = Lockwood) in which the protist is most abundant.

**Table S6** Taxonomic data of the 89 core taxa listed in the heatmap in Fig. 3. The table also indicates taxa that made up more than 1% of the total sequences, if they were enriched in either site, and if their V9 sequence matched that of a cultured isolate.

**Table S7** Data for the 103 protists that were isolated and cultured in this study.

**Figure S1.**
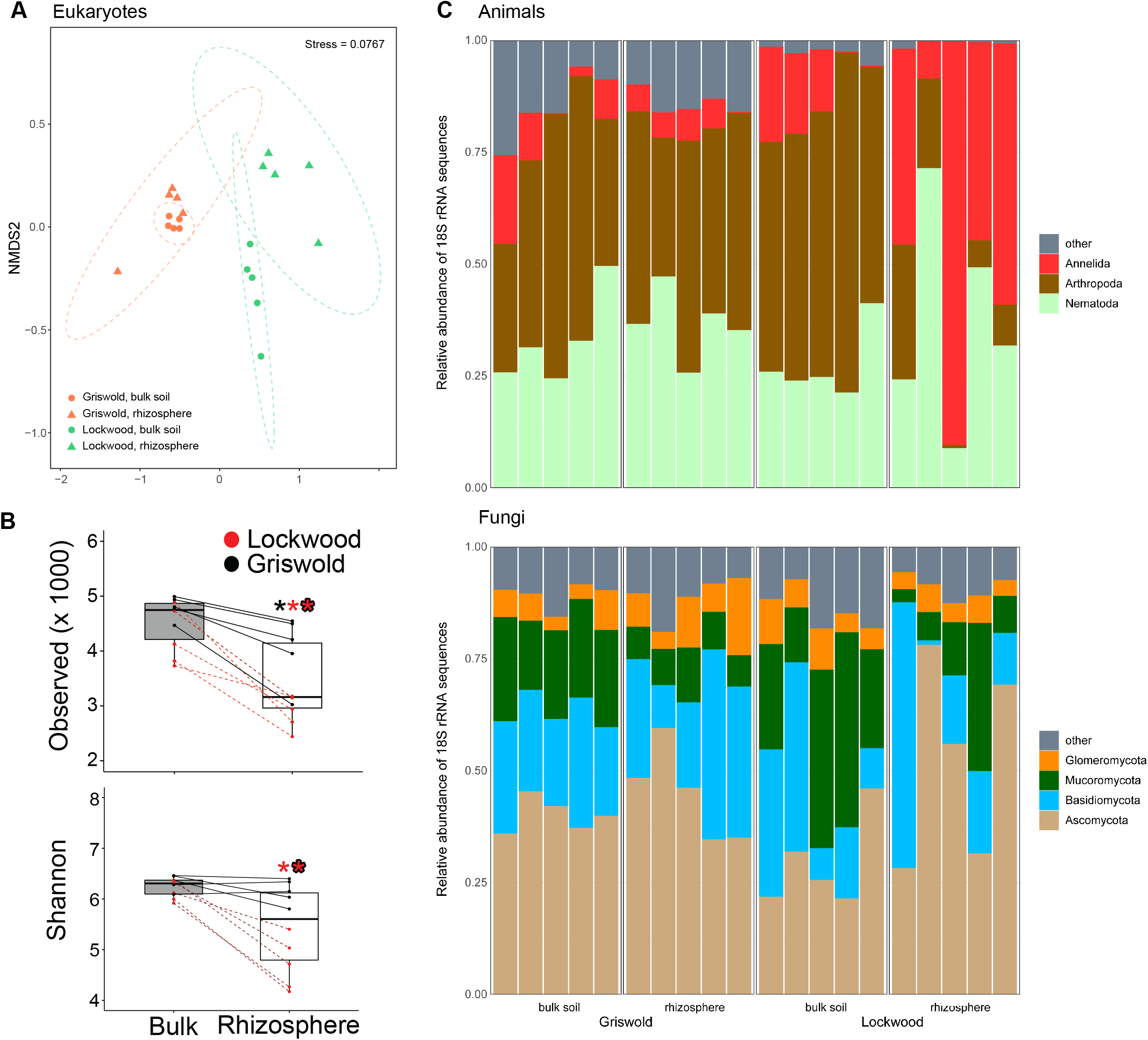
Community composition of total eukaryote samples. **A)** NMDS plot showing the clustering of samples of eukaryotic microbes (animals, fungi and protists). **B)** Diversity of the eukaryote and bacteria communities in the bulk soil and rhizosphere samples. Box plots indicate observed ASVs and Shannon Index. Lines between points in the bulk soil and rhizosphere samples indicate that the bulk soil and rhizosphere samples were obtained from the same plant root. **C)** Relative abundances of the most dominant phylum-level groups of animals and fungi. The category “other” represents the sum of all other taxonomic groups.

**Figure S2.**
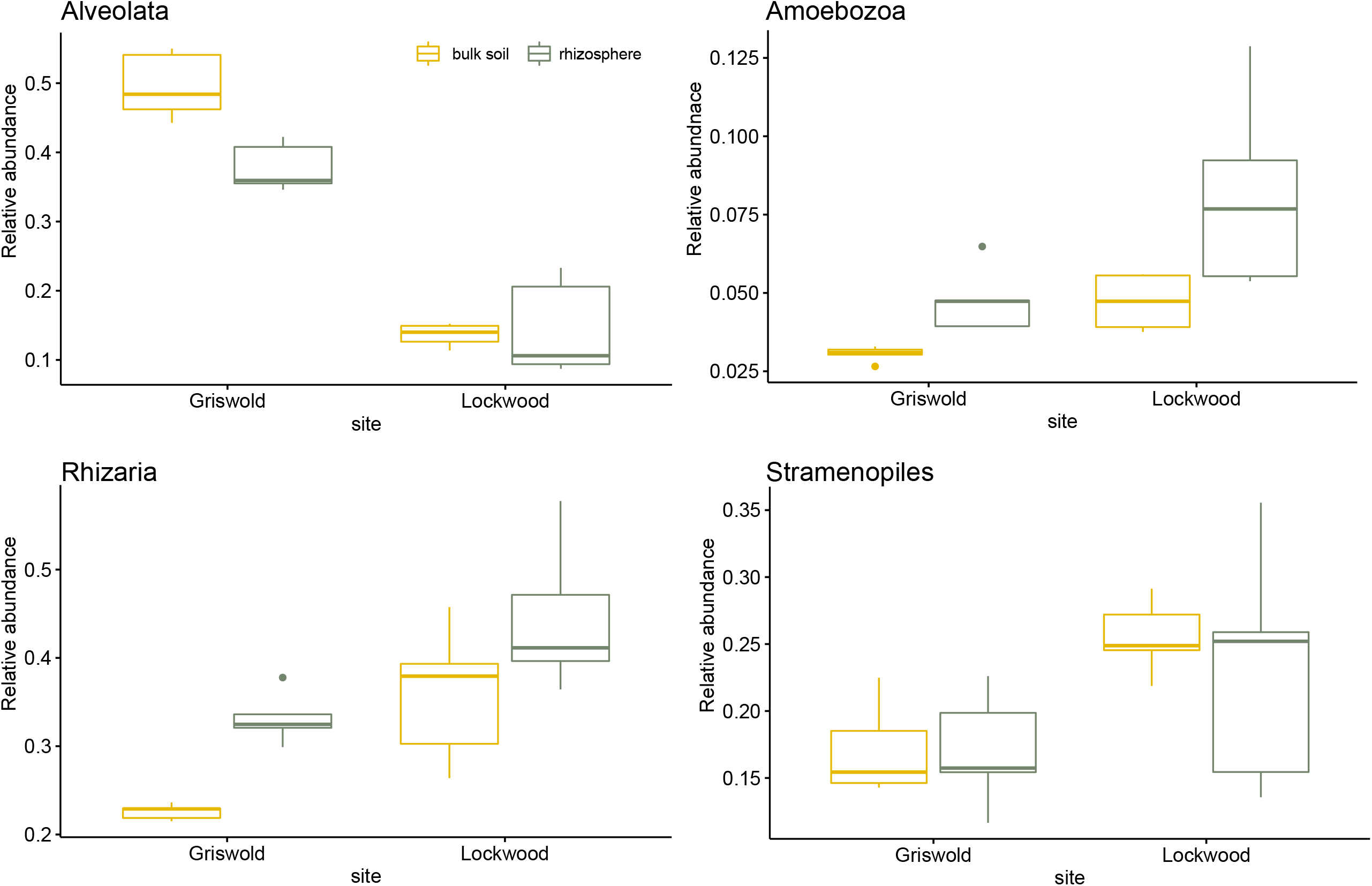
Boxplots showing the relative abundances of the four main supergroups associated with the rhizosphere samples across the different treatments.

**Table S1.** Site and sampling metadata for the maize plots at Griswold Research Station and Lockwood Farm.

| Site metadata |  |  |
| --- | --- | --- |
| Site | Griswold, CT, USA | Lockwood, CT, USA |
| Coordinates | 41.559708, -71.874195 | 41.407591, -72.907039 |
| Soil type and texture<sup>a</sup> | Windsor Loamy Sand | Cheshire Fine Sandy Loam |
| Cation Exchange Capacity<sup>b</sup> | 5 | 6.5 |
| K Saturation (%) | 4.5 | 5.6 |
| Mg Saturation (%) | 27.2 | 19.7 |
| Ca Saturation (%) | 46.4 | 52.8 |
| pH | 6.5 | 6.5 |
| Organic Matter (%) | 1.7 | 1.5 |
| Phosphorus (ppm) | 214 | 351 |
| Potassium (ppm) | 105 | 168 |
| Magnesium (ppm) | 185 | 174 |
| Calcium (ppm) | 618 | 913 |
| Previous season crop | vegetables | vegetables |
| Off-season plant composition (tilled under) | Cover crop: *Avena sativa* | Weeds: *Chenopodium album*, *Amaranthus retroflexus* |
| Sampling metadata |  |  |
| Pre-plant fertilizer (19-19-19) | 36 lbs/acre | 40 lbs/acre |
| Planting date | 15-May-19 | 17-May-19 |
| Sampling date | 10-Jul-19 | 12-Jul-19 |
| Air temp at sampling<sup>c</sup> | 90.8 | 87.9 |
| Growing Degree days from planting to sampling<sup>d</sup> | 915.6 | 1035.2 |
| Soil temp at sampling (°F) | 80.1 | 75.1 |
| Soil degree days from planting to sampling (°F) | 1348.8 | 938.7 |
| Cumulative rainfall from planting to sampling | 5.52" | 7.78" |
| Rainfall in the week prior to sampling | 0.0" | 0.37" |
| Percent water holding capacity at sampling<sup>e</sup> | 30.2 | 80.3 |

a. Soil type as determined by NRCS Web Soil Survey.
b. Soil nutrient, pH, and organic matter determination was determined on field samples taken at sampling time by Spectrum Analytic, Washington Court House, OH, USA.
c. Air and soil temperature and rain conditions were measured using a Hobo U30 station (Onset Computer Corporation, Bourne, MA 02532) placed close to each field. Soil measurements are at 20 cm depth.
d. Growing Degree Days for air and soil are accumulated degree days over 50° F.
e. Calculated as the volumetric water content on sampling day, as measured by Soil Moisture Smart Sensor (Onset Computer Corporation) divided by the maximum 2019 value at the site.

**Table S2.** Permanova and dispersion statistics for between-site and between-treatment comparison.

| Comparison | Taxa | PERMANOVA F-statistic |
| --- | --- | --- |
| Between Griswold and Lockwood | Bacteria | 9.61 |
|  | All eukaryotes | 8.01 |
|  | Animals | 5.23 |
|  | Fungi | 8.02 |
|  | Protists | 9.73 |
| Between bulk soil and rhizosphere in Griswold | Bacteria | 3.00 |
|  | All eukaryotes | 1.72 |
|  | Animals | 1.43 |
|  | Fungi | 1.40 |
|  | Protists | 2.44 |
| Between bulk soil and rhizosphere in Lockwood | Bacteria | 2.21 |
|  | All eukaryotes | 2.66 |
|  | Animals | 3.08 |
|  | Fungi | 2.00 |
|  | Protists | 2.47 |

son.
| PERMANOVA p-value | Dispersion F-statistic | Dispersion p-value |
| --- | --- | --- |
| **< 0.001** | 8.30 | **0.013** |
| **< 0.001** | 5.56 | **0.015** |
| **< 0.001** | 1.71 | 0.204 |
| **< 0.001** | 2.27 | 0.151 |
| **< 0.001** | 4.86 | **0.018** |
| **0.029** | 0.55 | 0.508 |
| **0.042** | 2.81 | 0.104 |
| 0.063 | 0.65 | 0.439 |
| 0.105 | 2.32 | 0.182 |
| **0.028** | 4.61 | **< 0.001** |
| **0.008** | 0.02 | 0.887 |
| **0.007** | 0.20 | 0.601 |
| **0.009** | 2.25 | 0.238 |
| **0.031** | 0.58 | 0.475 |
| **0.009** | 2.07 | 0.182 |

**Table S3.** Two-way analysis of variance (ANOVA) of the relative abundances of four most abundant protest supergroups across all samples. Site and soil compartment are the treatment factors.

|  | Df | Sum Sq | Mean Sq | F | P |
| --- | --- | --- | --- | --- | --- |
| site | 1 | 0.439 | 0.439 | 207.677 | **< 0.001** |
| soil_compartment | 1 | 0.015 | 0.015 | 7.010 | **0.018** |
| site x soil_compartment | 1 | 0.020 | 0.020 | 9.489 | **0.007** |
| Residuals | 16 | 0.034 | 0.002 |  |  |

**Amoebozoa**
|  | Df | Sum Sq | Mean Sq | F | P |
| --- | --- | --- | --- | --- | --- |
| site | 1 | 0.003 | 0.003 | 11.030 | **0.004** |
| soil_compartment | 1 | 0.003 | 0.003 | 11.531 | **0.004** |
| site x soil_compartment | 1 | 0.000 | 0.000 | 1.285 | 0.274 |
| Residuals | 16 | 0.005 | 0.000 |  |  |

**Rhizaria**
|  | Df | Sum Sq | Mean Sq | F | P |
| --- | --- | --- | --- | --- | --- |
| site | 1 | 0.076 | 0.076 | 21.849 | **< 0.001** |
| soil_compartment | 1 | 0.046 | 0.046 | 13.135 | **0.002** |
| site x soil_compartment | 1 | 0.001 | 0.001 | 0.159 | 0.695 |
| Residuals | 16 | 0.055 | 0.003 |  |  |

**Stramenopiles**
|  | Df | Sum Sq | Mean Sq | F | P |
| --- | --- | --- | --- | --- | --- |
| site | 1 | 0.026 | 0.026 | 9.030 | **0.008** |
| treatment_so | 1 | 0.001 | 0.001 | 0.247 | 0.626 |
| site:treatment_so | 1 | 0.001 | 0.001 | 0.243 | 0.629 |
| Residuals | 16 | 0.047 | 0.003 |  |  |

## References

Balczun, C. and Scheid, P.L. (2017) Free-living amoebae as hosts for and vectors of intracellular microorganisms with public health significance. Viruses 9: 65.

Bardgett, R.D. and Van Der Putten, W.H. (2014) Belowground biodiversity and ecosystem functioning. Nature 515: 505–511.

Bass, D. and del Campo, J. (2020) Microeukaryotes in animal and plant microbiomes:ecologies of disease? Eur J Protistol 76: 125719.

Bass, D., Howe, A.T., Mylnikov, A.P., Vickerman, K., Chao, E.E., Edwards Smallbone, J., et al. (2009) Phylogeny and classification of Cercomonadida (Protozoa, Cercozoa): Cercomonas, Eocercomonas, Paracercomonas, and Cavernomonas gen. nov. Protist 160: 483–521.

Berg, G. and Smalla, K. (2009) Plant species and soil type cooperatively shape the structure and function of microbial communities in the rhizosphere. FEMS Microbiol Ecol 68: 1–13.

Blandenier, Q., Seppey, C.V.W., Singer, D., Vlimant, M., Simon, A., Duckert, C., and Lara, E. (2017) Mycamoeba gemmipara nov. gen., nov. sp., the first cultured member of the environmental dermamoebidae clade LKM74 and its unusual life cycle. J Eukaryot Microbiol 64: 257–265.

Dumack, K., Feng, K., Flues, S., Sapp, M., Schreiter, S., Grosch, R., et al. (2020) What drives the assembly of plant-associated protist microbiomes? bioRxiv doi: 10.1101/2020.02.16.951384.

Flues, S., Bass, D., and Bonkowski, M. (2017) Grazing of leaf-associated Cercomonads (Protists: Rhizaria: Cercozoa) structures bacterial community composition and function. Environ Microbiol 19: 3297–3309.

Foissner, W. (1992) Estimating the species richness of soil protozoa using the “non flooded Petri dish method.” In Protocols in Protozoology. Lee, J.J. and T., S.A. (eds). Lawrence, KS: Allen Press, pp. 5–6.

Gao, Z., Karlsson, I., Geisen, S., Kowalchuk, G., and Jousset, A. (2019) Protists: puppet masters of the rhizosphere microbiome. Trends Plant Sci 24: 165–176.

Geisen, S. and Bonkowski, M. (2018) Methodological advances to study the diversity of soil protists and their functioning in soil food webs. Appl Soil Ecol 123: 328–333.

Guillou, L., Bachar, D., Audic, S., Bass, D., Berney, C., Bittner, L., et al. (2013) The Protist Ribosomal Reference database (PR^2^): a catalog of unicellular eukaryote Small Sub-Unit rRNA sequences with curated taxonomy. Nucleic Acids Res 41: 597–604.

Howe, A.T., Bass, D., Scoble, J.M., Lewis, R., Vickerman, K., Arndt, H., and Cavalier-Smith, T. (2011) Novel cultured protists identify deep-branching environmental DNA clades of Cercozoa: new genera Tremula, Micrometopion, Minimassisteria, Nudifila, Peregrinia. Protist 162: 332–372.

Howe, A.T., Bass, D., Vickerman, K., Chao, E.E., and Cavalier-Smith, T. (2009) Phylogeny, taxonomy, and astounding genetic diversity of Glissomonadida ord. nov., the dominant gliding zooflagellates in soil (Protozoa: Cercozoa). Protist 160: 159–189.

Jousset, A., Rochat, L., Scheu, S., Bonkowski, M., and Keel, C. (2010) Predator-prey chemical warfare determines the expression of biocontrol genes by rhizosphere-associated pseudomonas fluorescens. Appl Environ Microbiol 76: 5263–5268.

Keeling, P.J. and Campo, J. del (2017) Marine protists are not just big bacteria. Curr Biol 27: R541–R549.

Khojandi, N., Haselkorn, T.S., Eschbach, M.N., Naser, R.A., and DiSalvo, S. (2019) Intracellular Burkholderia symbionts induce extracellular secondary infections; driving diverse host outcomes that vary by genotype and environment. ISME J 13: 2068–2081.

Leach, J.E., Triplett, L.R., Argueso, C.T., and Trivedi, P. (2017) Communication in the phytobiome. Cell 169: 587–596.

Love, M.I., Huber, W., and Anders, S. (2014) Moderated estimation of fold change and dispersion for RNA-seq data with DESeq2. Genome Biol 15: 1–21.

Lowe, C.D., Martin, L.E., Roberts, E.C., Watts, P.C., Wootton, E.C., and Montagnes, D.J.S. (2011) Collection, isolation and culturing strategies for Oxyrrhis marina. J Plankton Res 33: 569–578.

Mahé, F., Vargas, C. De, Bass, D., Czech, L., Stamatakis, A., Lara, E., et al. (2017) Parasites dominate hyperdiverse soil protist communities in Neotropical rainforests. Nat Ecol Evol 1: 0091.

Marciano-Cabral, F. and Cabral, G. (2003) Acanthamoeba spp. as agents of disease in humans. Clin Microbiol Rev 16: 273–307.

McMurdie, P.J. and Holmes, S. (2013) Phyloseq: an R package for reproducible interactive analysis and graphics of microbiome census data. PLoS One 8: e61217.

McPherson, M.R., Wang, P., Marsh, E.L., Mitchell, R.B., and Schachtman, D.P. (2018) Isolation and analysis of microbial communities in soil, rhizosphere, and roots in perennial grass experiments. JoVE 137: e57932.

Nuccio, E.E., Anderson-Furgeson, J., Estera, K.Y., Pett-Ridge, J., De Valpine, P., Brodie, E.L., and Firestone, M.K. (2016) Climate and edaphic controllers influence rhizosphere community assembly for a wild annual grass. Ecology 97: 1307–1318.

Oliverio, A.M., Geisen, S., Delgado-Baquerizo, M., Maestre, F.T., Turner, B.L., and Fierer, N. (2020) The global-scale distributions of soil protists and their contributions to belowground systems. Sci Adv 6: 1–11.

Page, F.C. (1976) An Illustrated Key to Freshwater and Soil Amoebae with notes on Cultivation and Ecology, Ambleside, Cumbria, UK: Freshwater Biological Association, Scientific Publicatino No. 34.

Peiffer, J.A., Spor, A., Koren, O., Jin, Z., Tringe, S.G., Dangl, J.L., et al. (2013) Diversity and heritability of the maize rhizosphere microbiome under field conditions. Proc Natl Acad Sci U S A 110: 6548–6553.

Pérez, J., Munoz-Dorado, J., Brana, A.F., Shimkets, L.J., Sevillano, L., and Santamaría, R.I. (2011) Myxococcus xanthus induces actinorhodin overproduction and aerial mycelium formation by Streptomyces coelicolor. Microb Biotechnol 4: 175–183.

Philippot, L., Spor, A., Hénault, C., Bru, D., Bizouard, F., Jones, C.M., et al. (2013) Loss in microbial diversity affects nitrogen cycling in soil. ISME J 7: 1609–1619.

Quast, C., Pruesse, E., Yilmaz, P., Gerken, J., Schweer, T., Yarza, P., et al. (2013) The SILVA ribosomal RNA gene database project: improved data processing and web-based tools. Nucleic Acids Res 41: 590–596.

Rognes, T., Flouri, T., Nichols, B., Quince, C., and Mahé, F. (2016) VSEARCH: a versatile open source tool for metagenomics. PeerJ 4: e2584.

Rosenberg, K., Bertaux, J., Krome, K., Hartmann, A., Scheu, S., and Bonkowski, M. (2009) Soil amoebae rapidly change bacterial community composition in the rhizosphere of Arabidopsis thaliana. ISME J 3: 675–684.

Rossmann, M., Pérez-Jaramillo, J.E., Kavamura, V.N., Chiaramonte, J.B., Dumack, K., Fiore-Donno, A.M., et al. (2020) Multitrophic interactions in the rhizosphere microbiome of wheat: from bacteria and fungi to protists. FEMS Microbiol Ecol 96: 032.

Rubinstein, R.L., Kadilak, A.L., Cousens, V.C., Gage, D.J., and Shor, L.M. (2015) Protist-facilitated particle transport using emulated soil micromodels. Environ Sci Technol 49: 1384–1391.

Rüger, L., Feng, K., Dumack, K., Freudenthal, J., Chen, Y., Sun, R., et al. (2021) Assembly patterns of the rhizosphere microbiome along the longitudinal root axis of maize (Zea mays L.). Front Microbiol 12: 1–14.

Sapp, M., Ploch, S., Fiore-Donno, A.M., Bonkowski, M., and Rose, L.E. (2018) Protists are an integral part of the Arabidopsis thaliana microbiome. Environ Microbiol 20: 30–43.

Sasse, J., Martinoia, E., and Northen, T. (2018) Feed your friends: do plant exudates shape the root microbiome? Trends Plant Sci 23: 25–41.

Sato, T., Kuwahara, H., Fujita, K., Noda, S., Kihara, K., Yamada, A., et al. (2014) Intranuclear verrucomicrobial symbionts and evidence of lateral gene transfer to the host protist in the termite gut. ISME J 8: 1008–1019.

Schloss, P.D., Westcott, S.L., Ryabin, T., Hall, J.R., Hartmann, M., Hollister, E.B., et al. (2009) Introducing mothur: open-source, platform-independent, community-supported software for describing and comparing microbial communities. Appl Environ Microbiol 75: 7537–7541.

Shetty, S.A. and Lahti, L. (2018) microbiomeutilities: An R package for utilities to guide in-depth marker gene amplicon data analysis. R Packag Version 2128.

Sibbald, S.J. and Archibald, J.M. (2017) More protist genomes needed. Nat Ecol Evol 1: 1–3.

Sun, A., Jiao, X., Chen, Q., Trivedi, P., Li, Z., Li, F., et al. (2021) Fertilization alters protistan consumers and parasites in crop-associated microbiomes. Environ Microbiol early view.

Taerum, S.J., Steven, B., Gage, D.J., and Triplett, L.R. (2020) Validation of a PNA clamping method for reducing host DNA amplification and increasing eukaryotic diversity in rhizosphere microbiome studies. Phytobiomes J 4: 291–302.

Takenouchi, Y., Iwasaki, K., and Murase, J. (2016) Response of the protistan community of a rice field soil to different oxygen tensions. FEMS Microbiol Ecol 92: 1–8.

Trivedi, P., Leach, J.E., Tringe, S.G., Sa, T., and Singh, B.K. (2020) Plant–microbiome interactions: from community assembly to plant health. Nat Rev Microbiol 18: 607–621.

Vaulot, D., Mahé, F., and Geisen, S. (2020) pr2-primer: an 18S rRNA primer database for protists. bioRxiv doi: 10.1101/2021.01.04.425170.

Wang, Q., Garrity, G.M., Tiedje, J.M., and Cole, J.R. (2007) Naïve Bayesian classifier for rapid assignment of rRNA sequences into the new bacterial taxonomy. Appl Environ Microbiol 73: 5261–5267.

Weidner, S., Latz, E., Agaras, B., Valverde, C., and Jousset, A. (2017) Protozoa stimulate the plant beneficial activity of rhizospheric pseudomonads. Plant Soil 410: 509–515.

Xiong, W., Song, Y., Yang, K., Gu, Y., Wei, Z., Kowalchuk, G.A., et al. (2020) Rhizosphere protists are key determinants of plant health. Microbiome 8: 27.

Zhang, L. and Lueders, T. (2017) Micropredator niche differentiation between bulk soil and rhizosphere of an agricultural soil depends on bacterial prey. FEMS Microbiol Ecol 93: 1– 11.

Zou, S., Zhang, Q., Zhang, X., Dupuy, C., and Gong, J. (2020) Environmental factors and pollution stresses select bacterial populations in association with protists. Front Mar Sci 7: 1–19.

