## Supplementary material for "A dual sequence and culture-based survey of maize rhizosphere protists reveals dominant, plant-enriched, and culturable community members": Table S4

**Table S4.** Relative abundances of the top five protist families in each supergroup within sites and plant compartments. Taxa names are as designated by families representing greater than a mean of 5% of protist reads are shown in bold. The family in the top five most abundant in each rhizosphere, but not in the bulk soils, is underlined.

|  |  | **Griswold** | | | | **Lockwood** | | | |
| --- | --- | --- | --- | --- | --- | --- | --- | --- | --- |
|  |  | **Bulk** | | **Rhizosphere** | | **Bulk** | | **Rhizosphere** | |
|  | Rank | Family | Relative abundance^a^ | Family | Relative abundance | Family | Relative abundance | Family | Relative abundance |
| **Alveolata** | 1 | **Actinocephalidae** | **38.48±5.12%** | **Actinocephalidae** | **27.75±3.99%** | Colpodida (uncl.) | 3.25±1.03% | **Colpodida (uncl.)** | **8.18±6.99%** |
|  | 2 | Hypotrichia (uncl.)^b^ | 2.02±0.63% | Hypotrichia (uncl.) | 2.35±0.45% | Actinocephalidae | 2.43±0.77% | Hypotrichia (uncl.) | 1.96±1.18% |
|  | 3 | Colpodida (uncl.) | 1.68±0.42% | Colpodida (uncl.) | 1.77±0.61% | Hypotrichia (uncl.) | 2.17±0.94% | Actinocephalidae | 1.51±0.7% |
|  | 4 | Ciliophora (uncl.) | 1.67±0.4% | Litostomatea (uncl.) | 1.36±0.3% | Colpodea X (uncl.) | 1.28±0.33% | Colpodea X (uncl.) | 0.69±0.45% |
|  | 5 | Litostomatea (uncl.) | 1.64±0.74% | Colpodea X (uncl.) | 1.03±0.4% | Ciliophora (uncl.) | 0.98±0.34% | Platyophryida | 0.44±0.31% |
| **Rhizaria** | 1 | Cercomonadidae | 4.81±0.65% | **Cercomonadidae** | **7.92±4.03%** | **Cercomonadidae** | **7.46±1.87%** | **Cercomonadidae** | **11.93±1.5%** |
|  | 2 | Cryomonadida (uncl.) | 3.02±0.7% | **Paracercomonadidae** | **5.82±1.12%** | **Rhogostoma-lineage** | **5.52±2.14%** | **Allapsidae** | **9.9±3.05%** |
|  | 3 | Rhogostoma-lineage | 2.9±0.97% | Allapsidae | 4.46±2.96% | Cryomonadida (uncl.) | 4.51±2.08% | **Glissomonadida (**uncl.) | **6.18±5.01%** |
|  | 4 | Paracercomonadidae | 2.17±0.27% | Cryomonadida (uncl.) | 3.57±1.24% | Glissomonadida (uncl.) | 2.7±1.32% | Cryomonadida (uncl.) | 3.59±2.5% |
|  | 5 | Leptophryidae | 1.59±0.72% | Rhogostoma-lineage | 3.13±0.67% | Paracercomonadidae | 2.36±0.75% | Paracercomonadidae | 2.85±1.86% |
| **Stramenopile** | 1 | Chrysophyceae Clade-C | 3.71±1.27% | **Peronosporales (uncl.)** | **6.27±5.07%** | **Chrysophyceae X (**uncl.) | **5.97±1.55%** | **Chrysophyceae X (**uncl.) | **12.6±8.18%** |
|  | 2 | Peronosporales (uncl.) | 3.54±2.74% | Chrysophyceae Clade-C | 2.75±0.25% | Peronosporales (uncl.) | 4.66±1.42% | Peronosporales (uncl.) | 2.93±1.4% |
|  | 3 | Chrysophyceae X (uncl.) | 2.07±0.37% | Hyphochytriaceae | 1.63±0.48% | Chrysophyceae Clade-C | 4.36±2.13% | Chrysophyceae Clade-C | 1.88±0.98% |
|  | 4 | Hyphochytriaceae | 1.95±0.19% | Chrysophyceae X (uncl.) | 1.5±0.43% | Hyphochytriaceae | 2.31±0.63% | Ochrophyta (uncl.) | 1.62±0.54% |
|  | 5 | Ochrophyta (uncl.) | 1.28±0.16% | Oomycota X (uncl.) | 1.09±0.82% | Ochrophyta (uncl.) | 2.22±0.52% | Hyphochytriaceae | 1.02±0.39% |
| **Amoebozoa** | 1 | Acanthamoebidae | 0.93±0.16% | Acanthamoebidae | 0.96±0.32% | Acanthamoebidae | 1.31±0.35% | Vermamoebidae | 1.41±0.77% |
|  | 2 | Filamoebidae | 0.45±0.04% | Filamoebidae | 0.88±0.43% | Filamoebidae | 0.9±0.31% | Filamoebidae | 1.36±0.51% |
|  | 3 | Leptomyxidae | 0.36±0.15% | Mb5C-lineage | 0.51±0.34% | LKM74-lineage | 0.32±0.11% | Acanthamoebidae | 1.3±0.43% |
|  | 4 | Tubulinea (uncl.) | 0.19±0.11% | LKM74-lineage | 0.39±0.08% | Tubulinea (uncl.) | 0.28±0.13% | LKM74-lineage | 1.22±0.38% |
|  | 5 | Mb5C-lineage | 0.18±0.08% | Tubulinea (uncl.) | 0.34±0.25% | Leptomyxidae | 0.27±0.15% | Tubulinea (uncl.) | 1.07±1.25% |

a. Mean percentage of all reads in each sample represented by each family, plus or minus the standard deviation (N=5).

b. Uncl. = belongs to an unclassified family within that class or order
